# CAPRI: Efficient Inference of Cancer Progression Models from Cross-sectional Data

**DOI:** 10.1101/008110

**Authors:** Daniele Ramazzotti, Giulio Caravagna, Loes Olde Loohuis, Alex Graudenzi, Ilya Korsunsky, Giancarlo Mauri, Marco Antoniotti, Bud Mishra

## Abstract

We devise a novel inference algorithm to effectively solve the *cancer progression model reconstruction* problem. Our empirical analysis of the accuracy and convergence rate of our algorithm, *CAncer PRogression Inference* (CAPRI), shows that it outperforms the state-of-the-art algorithms addressing similar problems.

**Motivation:** Several cancer-related genomic data have become available (e.g., *The Cancer Genome Atlas*, TCGA) typically involving hundreds of patients. At present, most of these data are aggregated in a *cross-sectional* fashion providing all measurements at the time of diagnosis.

Our goal is to infer cancer “progression” models from such data. These models are represented as directed acyclic graphs (DAGs) of collections of “*selectivity*” relations, where a mutation in a gene *A* “selects” for a later mutation in a gene *B*. Gaining insight into the structure of such progressions has the potential to improve both the stratification of patients and personalized therapy choices.

**Results:** The CAPRI algorithm relies on a scoring method based on a *probabilistic theory* developed by Suppes, coupled with *bootstrap* and *maximum likelihood* inference. The resulting algorithm is efficient, achieves high accuracy, and has good complexity, also, in terms of convergence properties. CAPRI performs especially well in the presence of noise in the data, and with limited sample sizes. Moreover CAPRI, in contrast to other approaches, robustly reconstructs different types of confluent trajectories despite irregularities in the data.

We also report on an ongoing investigation using CAPRI to study *atypical Chronic Myeloid Leukemia*, in which we uncovered non trivial selectivity relations and exclusivity patterns among key genomic events.

**Availability:** CAPRI is part of the *TRanslational ONCOlogy* R package and is freely available on the web at: http://bimib.disco.unimib.it/index.php/Tronco

## 1 Introduction

Analysis and interpretation of the fast-growing biological data sets that are currently being curated from laboratories all over the world require sophisticated computational and statistical methods.

Motivated by the availability of genetic patient data, we focus on the problem of *reconstructing progression models* of cancer. In particular, we aim to infer the plausible sequences of *genomic alterations* that, by a process of *accumulation*, selectively make a tumor fitter to survive, expand and diffuse (i.e., metastasize). Along the trajectories of progression, a tumor (monotonically) acquires or “activates” mutations in the genome, which, in turn, produce progressively more “viable” clonal subpopulations over the so-called *cancer evolutionary landscape* (cfr., [1, 2, 3]).

Knowledge of such progression models is very important for drug development and in therapeutic decisions. For example, it has been known that for the same cancer type, patients in different stages of different progressions respond differently to different treatments.

Several datasets are currently available that aggregate diverse cancer-patient data and report in-depth mutational profiles, including e.g., structural changes (e.g., inversions, translocations, copynumber variations) or somatic mutations (e.g., point mutations, insertions, deletions, etc.). An example of such a dataset is *The Cancer Genome Atlas* (TCGA) (cfr.,[ 4])). These data, by their very nature, only give a snapshot of a given tumor sample, mostly from biopsies of untreated tumor samples at the time of diagnoses. It still remains impractical to track the tumor progression in any single patient over time, thus limiting most analysis methods to work with *cross-sectional* data^1^.

To rephrase, we focus on the problem of *cancer progression models reconstruction from cross-sectional data*. The problem is not new and, to the best of our knowledge, two threads of research starting in the late 90’s have addressed it. The first category of works examined mostly gene-expression data to reconstruct the temporal ordering of samples (cfr., [5, 6]). The second category of works looked at inferring cancer progression models of increasing model-complexity, starting from the simplest tree models (cfr. [7]) to more complex graph models (cfr., [8]); see the next subsection for an overview of the state of the art. Building on our previous work described in [9] we present a novel and comprehensive algorithm of the second category that addresses this problem.

The new algorithm proposed here is called *CAncer PRogression Inference* (CAPRI) and is part of the *TRanslational ONCOlogy* (TRONCO) package (cfr., [10]). Starting from cross-sectional genomic data, CAPRI reconstructs a probabilistic progression model by inferring “selectivity relations”, where a mutation in a gene *A* “selects” for a later mutation in a gene *B*. These relations are depicted in a combinatorial graph and resemble the way a mutation exploits its “*selective advantage*” to allow its host cells to expand clonally. Among other things, a selectivity relation implies a putatively invariant temporal structure among the genomic alterations (i.e., *events*) in a specific cancer type. In addition, these relations are expected to also imply “probability raising” for a pair of events in the following sense: Namely, a selectivity relation between a pair of events here signifies that the presence of the earlier genomic alteration (i.e., the *upstream event*) that is advantageous in a Darwinian competition scenario increases the probability with which a subsequent advantageous genomic alteration (i.e., the *downstream event*) appears in the clonal evolution of the tumor. Thus the selectivity relation captures the effects of the evolutionary processes, and not just correlations among the events and imputed clocks associated with them. As an example, we show in (Figure 1) the selectivity relation connecting a mutation of egfr to the mutation of CDK.

Consequently, an inferred selectivity relation suggests mutational profiles in which certain samples (early-stage patients) display specific alterations only (e.g., the alteration characterizing the beginning of the progression), while certain other samples (e.g., late-stage patients) display a superset subsuming the early mutations (as well as alterations that occur subsequently in the progression).

**Figure 1:**
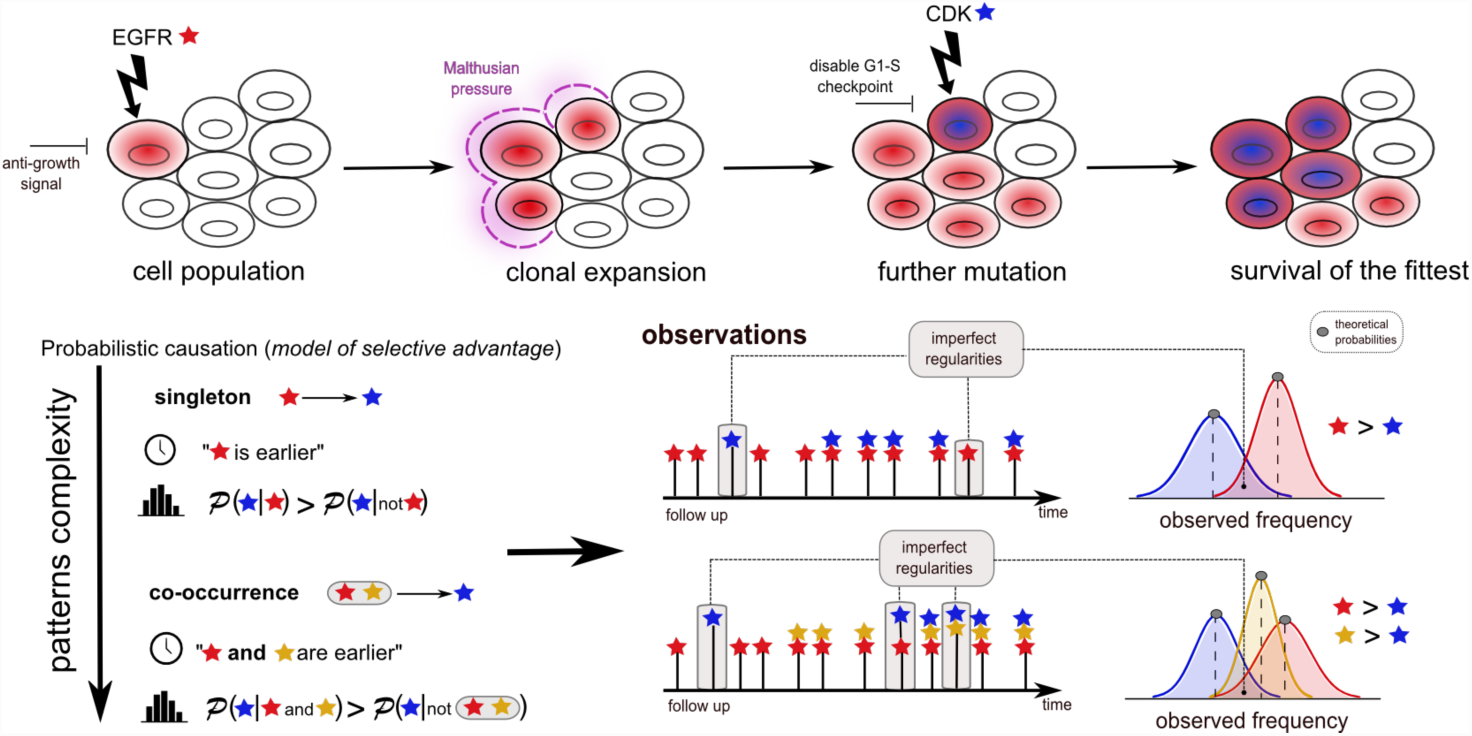
Selectivity Relation in Tumor Evolution. The *CAncer PRogression Inference* (CAPRI) algorithm examines cancer patients’ genomic cross-sectional data to determine relationships among genomic alterations (e.g., somatic mutations, copy-number variations, etc.) that modulate the somatic evolution of a tumor. When CAPRI concludes that aberration *a* (say, an egfr mutation) “selects for” aberration *b* (say, a cdk mutation), such relations can be rigorously expressed using Suppes’ conditions, which postulates that if *a* selects *b*, then *a* occurs before *b* (*temporal priority*) and occurrences of *a* raises the probability of emergence of *b* (*probability raising*). Moreover, CAPRI is capable of reconstructing relations among more complex boolean combination of events, as shown in the bottom panel and discussed in the Approach section.

Various kinds of genomic aberrations are suitable as input data, and include somatic point/indel mutations, copy-number alterations, etc., provided that they are *persistent*, i.e., once an alteration is acquired no other genomic event can restore the cell to the non-mutated (i.e., *wild type*) condition^2^.

The selectivity relations that CAPRI reconstructs are ranked and subsequently further refined by means of a hybrid algorithm, which reasons over time, mechanism and chance, as follows. CAPRI’s overall scoring methods combine topological constraints grounded on Patrick Suppes’ conditions of probabilistic causation (see e.g., [12]), with a *maximum likelihood-fit* procedure (cfr., [13]) and derives much of its statistical power from the application of *bootstrap* procedures (see e.g., [14]). CAPRI returns a *graphical model* of a complex selectivity relation among events which captures the essential aspects of cancer evolution: branches, confluences and independent progressions. In the specific case of confluences, CAPRI’s ability to infer them is related to the complexity of the “patterns” they exhibit, expressed in a logical fashion. As pointed out by other approaches (cfr., [15]), this strategy requires trading off complexity for expressivity of the inferred models, and results in two execution modes for the algorithm: supervised and unsupervised, which we discuss in details in Sections 2 and 3.

In Section 3 (Methods) we show that CAPRI enjoys a set of attractive properties in terms of its complexity, soundness and expressivity, even in the presence of uniform *noise* in the input data – e.g., due to *genetic heterogeneity* and experimental errors. Although many other approaches enjoy similar asymptotic properties, we show that CAPRI can compute accurate results with surprisingly small sample sizes (cfr., Section 4). Moreover, to the best of our knowledge, based on extensive synthetic data simulations, CAPRI outperforms all the competing procedures with respect to all desirable performance metrics. We conclude by showing an application of CAPRI to reconstruct a progression model for *atypical Chronic Myeloid Leukemia* (aCML) using a recent exome sequencing dataset, first presented in [16].

### 1.1 State of the Art

For an extensive review on *cancer progression model reconstruction* we refer to the recent survey by [17]. In brief, progression models for cancer have been studied starting with the seminal work of [18] where, for the first time, cancer progression was described in terms of a directed path by assuming the existence of a unique and most likely temporal order of genetic mutations. [18] manually created a (colorectal) cancer progression from a genetic and clinical point of view. More rigorous and complex algorithmic and statistical automated approaches have appeared subsequently. As stated already, the earliest thread of research simply sought more generic progression models that could assume tree-like structures. The *oncogenetic tree model* captured evolutionary branches of mutations (cfr., [7, 19]) by optimizing a *correlation*-based score. Another popular approach to reconstruct tree structures appears in [20]. Other general Markov chain models such as, e.g., [21] reconstruct more flexible probabilistic networks, despite a computationally expensive parameter estimation. In [9], we introduced an algorithm called *CAncer PRogression Extraction with Single Edges* (CAPRESE), which, based on its extensive empirical analysis, may be deemed as the current state-of-the-art algorithm for the inference of tree models of cancer progression. It is based on a shrinkage-like statistical estimation, grounded in a general theoretical framework, which we extend further in this paper. Other results that extend tree representations of cancer evolution exploit mixture tree models, i.e., multiple oncogenetic trees, each of which can independently result in cancer development (cfr., [22]). In general, all these methods are capable of modeling diverging temporal orderings of events in terms of branches, although the possibility of converging evolutionary paths is precluded.

To overcome this limitation, the most recent approaches tends to adopt Bayesian graphical models, i.e., Bayesian Networks (BN). In the literature, there have been two initial families of methods aimed at inferring the structure of a BN from data (cfr., [13]). The first class of models seeks to explicitly capture all the conditional independence relations encoded in the edges and will be referred to as *structural approaches*; the methods in this family are inspired by the work on causal theories by Judea Pearl (cfr., [23, 24, 25, 26]). The second class – *likelihood approaches* – seeks a model that maximizes the likelihood of the data (cfr., [27, 28, 29]).

A more recent *hybrid approach* to learn a BN which combines the two families above by (*i*) constraining the search space of the valid solutions and, then, (*ii*) fitting the model with likelihood maximization (see [15, 8, 30]). A further technique to reconstruct progression models from cross-sectional data was introduced in [31], in which the transition probabilities between genotypes are inferred by defining a Moran process that describes the evolutionary dynamics of mutation accumulation. In [32] this methodology was extended to account for pathway-based phenotypic alterations.

## 2 Approach

In what follows, we denote with 𝒫(·) and 𝒫(· *|* ·) the observed marginal and conditional probability of an event, whose complement is denoted with the diacritical mark · (macron).

### A probabilistic model of selective advantage

Central to CAPRI’s score function is Suppes’ notion of *probabilistic causation* (cfr., [12]), which can be stated in the following terms: a selectivity relation^3^ among two observables *i* and *j* if (1) *i* occurs earlier than *j* – *temporal priority* (tp) – and (2) if the probability of observing *i* raises the probability of observing *j*, i.e., 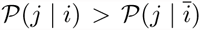 – *probability raising* (pr). The definition of probability raising subsumes positive statistical dependency and mutuality (see, e.g., [9]). Note that the resulting relation (also, called prima facie causality) is purely observational and remains agnostic to the possible mechanistic cause-effect relation involving *i* and *j*.

While Suppes’ definition of probabilistic causation has known limitations in the context of general causality theory (see discussions in, e.g., [33, 34]), in the context of cancer evolution, this relation appropriately describes various features of *selective advantage* in somatic alterations that accumulate as tumor progresses.

Thus, in our framework, we implement the temporal priority among events – condition (1) – as 𝒫 (*i*) > 𝒫 (*j*), because it is intuitively sound to assume that the (cumulative) genomic events occurring earlier are the ones present in higher frequency in a dataset. In addition, condition (2) is implemented as is, that is by requiring that for each pair of observables *i* and *j* directly connected, 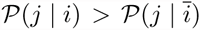 is verified. Taken together, these conditions gives rise to a natural ordering relation among events, written “*i* ▹ *j*” and read as “*i* has a selective influence on *j*.” This relation is a *necessary* but *not sufficient* condition to capture the notion of selective advantage, and additional constraints need to be imposed to filter spurious relations. Spurious correlations are both intrinsic to the definition (e.g., if *i* ▹ *j* ▹ *w* then also *i* ▹ *w*, which could be spurious) and to the model we aim at inferring, because data is finite as well as corrupted by noise.

Building on this framework, we devise inference algorithms that capture the essential aspects of heterogeneous cancer progressions: *branching*, *independence* and *convergence* – all combining in a progression model.

### Progression patterns

The complexity of cancer requires modeling multiple non-trivial *patterns* of its progression: for a specific event, a pattern is defined as a specific combination of the closest upstream events that confers a selective advantage.

As an example, imagine a clonal subpopulation becoming fit – thus enjoying expansion and selection – once it acquires a mutation of gene *c*, provided it also has previously acquired a mutation in a gene in the upstream *a*/*b* pathway. In terms of progression, we would like to capture the trajectories: {*a,* ¬*b* }, {¬*a, b* } and {*a, b*} precedes *c* (where ¬ denotes the absence of an event in the gene).

To establish this analysis formally, we augment our model of selection in a tumor with a language built from simple propositional logic formulas using the usual Boolean connectives: namely, “and” (⋀), “or” (⋁) and “xor” (⨁). These patterns can be described by formulæ in a propositional logical language, which can be rendered in *Conjunctive Normal Form* (CNF). A CNF formula *φ* has the following syntax: *φ* = ***𝒸***_1_ ⋀ *…* ⋀ ***𝒸***_*n*_, where each ***𝒸***_*i*_ is a *disjunctive clause* ***𝒸***_*i*_ = *c*_*i,*1_ ⋁ *…* ⋁ *c*_*i,k*_ over a set of literals, each literal representing an event or its negation. Given this (rather obvious) pattern representation, we write the conditions for *selectivity with patterns* as

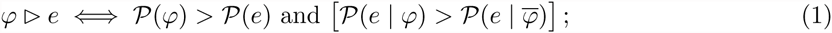

with respect to the example above, patterns ^4^ could be *a* ⋁ *b* ▹ *c* and *a* ⊕ *b* ▹ *c*

In our framework the problem of reconstructing a probabilistic graphical model of progression reduces to the following: for each input event *e*, assess a *set of selectivity patterns* {*φ*_1_ ▹ *e, …, φ*_*k*_ ▹ *e*}, filter the spurious ones, and combine the rest in a *direct acyclic graph* (DAG)^5^, augmented with logical symbols. Notice that while we broke down the progression extraction into a series of sub-tasks, the problem remains complex: patterns are unknown, potentially spurious, and exponential in formula size; data is noisy; patterns must allow for “imperfect regularities”, rather than being strict^6^. To summarize, in our setting we can model complex progression trajectories with branches (i.e., events involved in various patterns), independent progressions (i.e., events without common ancestors) and convergence (via CNF formulas). The framework we introduce here is highly versatile, and to the best of our knowledge, it infers and checks more complex claims than any cancer progression algorithms described thus far (cfr.,[7, 8, 9]).

**Figure 2:**
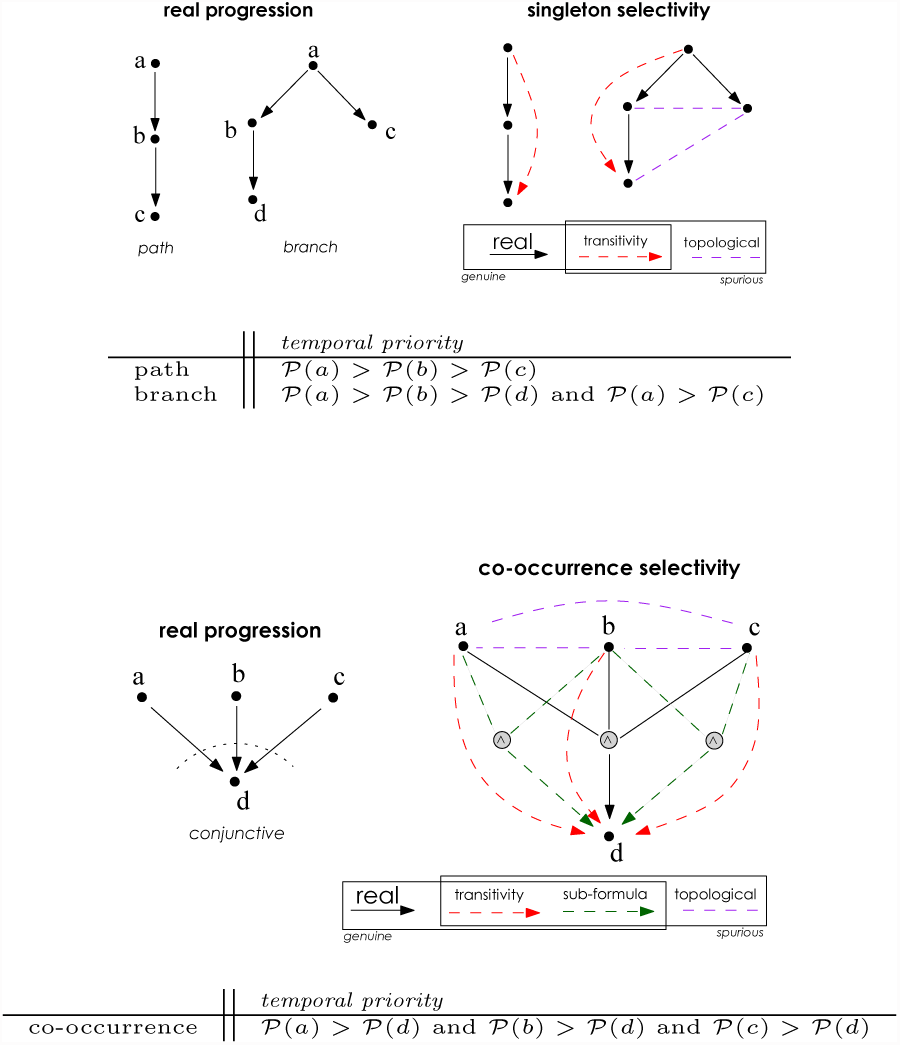
Singleton and Co-occurrence Selectivity Patterns. Examples of patterns that CAPRI can automatically extract without prior hypotheses. *(Top):* A linear path and branching model (left) and corresponding singleton selectivity patterns with infinite sample size (right). All the genuine connections are shown (red and black, directed by the temporal priority), as well as edges (purple, undirected) which might be suggested by the topology (or observations, if data were finite). *(Bottom):* Example of conjunctive model (*a and b and c*). The co-occurrence selectivity pattern is shown, with all true patterns and infinite sample size. The topology is augmented by logical connectives; green arrows are spurious patterns emerging from the structure of the true pattern *a* ⋀ *b* ⋀ *c* ▹ *d*.

**Figure 3:**
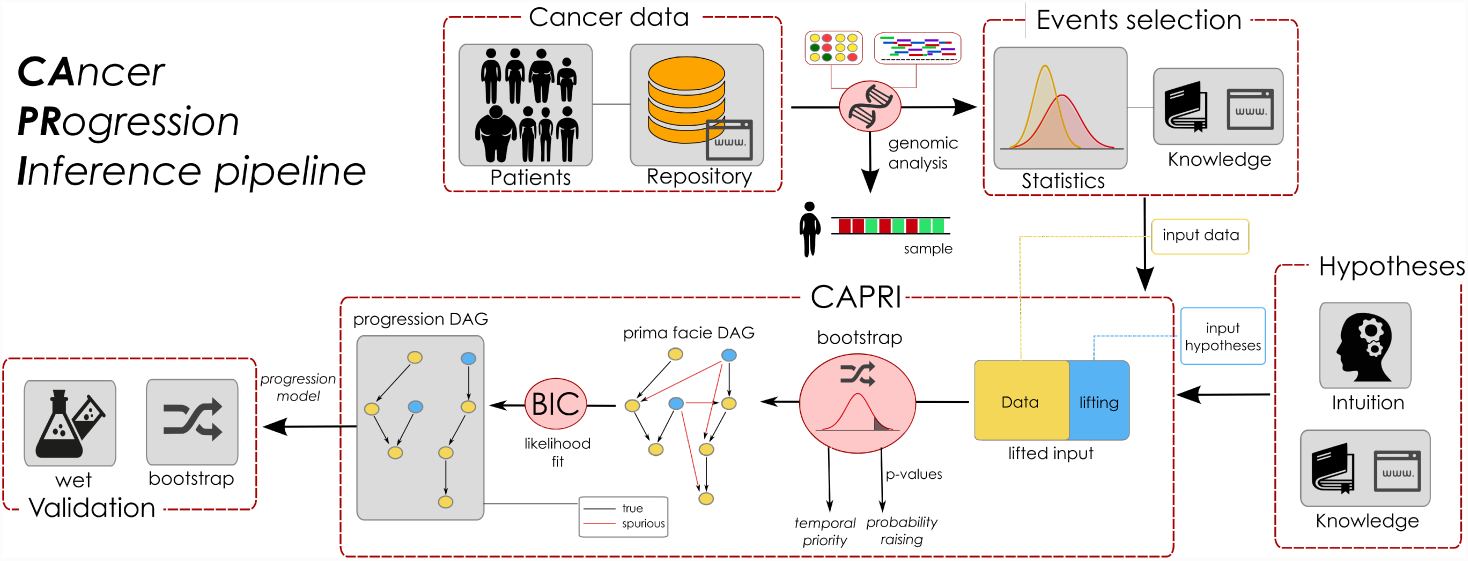
Data processing pipeline for cancer progression inference. We sketch a pipeline to best exploit CAPRI’s ability to extract cancer progression models from cross-sectional data. Initially, one collects *experimental data* (which could be accessible through publicly available repositories such as TCGA) and performs *genomic analyses* to derive profiles of, e.g., somatic mutations or Copy-Number Variations for each patient. Then, statistical analysis and biological priors are used to select events relevant to the progression and imputable by CAPRI - e.g., *driver mutations*. To exploit CAPRI’s supervised execution mode (see Methods) one can use further statistics and priors to generate *patterns of selective advantage* -, e.g, hypotheses of mutual exclusivity. CAPRI can extract a progression model from these data and assess various *confidence* measures on its constituting relations - e.g., (non-)parametric bootstrap and hypergeometric testing. *Experimental validation* concludes the pipeline.

## 3 Methods

Building on the framework described in the previous section, we now describe the implementation of CAPRI’s building blocks. Notice that, in general, the inference of cancer progression models requires a complex *data processing pipeline*, as summarized in Figure 3; its architecture optimally exploits CAPRI’s efficiency.

### Assumptions

CAPRI relies on the following assumptions: *i*) Every pattern is expressible as a propositional CNF formula; *ii*) All events are persistent, i.e., an acquired mutation cannot disappear; *iii*) All relevant events in tumor progression are observable, with the observations describing the progressive phenomenon in an essential manner (i.e., *closed world* assumption, in which all events ‘driving’ the progression are detectable); *iv*) All the events have non-degenerate observed probability in (0, 1); *v*) All events are distinguishable, in the following sense: input alterations produce different profiles across input samples. Assumptions *i*-*ii*) relate to the framework derived in previous section, while *iii*) imposes an onerous burden on the experimentalists, who must select the *relevant* genomic events to model^7^. Assumption *iv*) relates instead to the statistical distinguishability of the input events (see the next section on CAPRI’s Data Input)‥

#### Trading Complexity for Expressivity

To automatically extract the patterns that underly a progression model, one may try to adopt a brute-force method of enumerating and testing all possibilities. This strategy is computationally intractable, however, since the number of (distinct) (sub)formulæ grows exponentially with the number of events included in the model. Therefore, we need to exploit certain properties of the ▹ relation whenever possible, and trade expressivity for complexity in other cases, as explained below.

Note that *singleton* and *co-occurrence* (⋀) types of patterns are amenable to *compositional reasoning* : if *i*_1_⋀ *…* ⋀*i*_*k*_ ▹ *j* then, for any *p* = 1*, …, k*, *i*_*p*_ ▹ *j*. This observation leads to the following straightforward strategy of evaluating every conjunctive (and henceforth singleton) relation using a pairwise-test for the selectivity relation (see Figure 2).

Unfortunately, it is easy to see that this reasoning fails to generalize for CNF patterns: e.g., when the pattern contains disjunctive operators (⋁). As an example, consider pattern *a* ⋁ *b* ▹ *c*, in a cancer where {*a,* ¬*b*} progression to *c* is more prevalent than {¬*a, b*} and {*a, b*}. In this case, considering sub-formulas only we might find *a* ▹ *c* but miss *b* ▹ *c* because the probability of mutated *b* is smaller than that of *c*, thus invalidating condition (1) of relation ▹. Notice that in extreme situations, when the data is very noisy, the algorithm may even “invert” the selectivity relation to *c* ▹ *b*.

This difficulty is not a peculiarity of our framework, but rather intrinsic to the problem of extracting complex “causal networks” (cfr., [23, 24, 34]). To handle this situation, CAPRI adapts a strategy that trades complexity for expressivity: the resulting inference procedure, Algorithm 1, can be executed in two modes: unsupervised and supervised. In the former, inferred patterns of confluent progressions are constrained to co-occurrence types of relations, in the latter CAPRI can test more complex patterns, i.e., disjunctive or “mutual exclusive” ones, provided they are given as prior hypotheses. In both cases, CAPRI’s complexity – studied in next sections – is quadratic both in the number of events and hypotheses.

#### Data Input (Step 1)

CAPRI (cfr., Algorithm 1) requires an input set *G* of *n* events, i.e., genomic alterations, and *m* cross-sectional samples, represented as a dataset in an *m* × *n* binary matrix *D*, in which an entry *D*_*i,j*_ = 1 if the event *j* was observed in sample *i*, and 0 otherwise. Assumption *iv*) is satisfied when all columns in *D* differ - i.e., the alteration profiles yield different observations.

Optionally, a set of *k* input hypotheses φ = *{φ*_1_ ▹ *e*_1_*, …, φ*_*k*_ ▹ *e*_*k*_*}*, where each *φ*_*i*_ is a well-formed^8^ CNF formula. Note that we advise that the algorithm be used in the following regime^9^: *k* + *n* ≪ *m*.

#### Data Preprocessing (Lifting, step 2)

When input hypotheses are provided (e.g., by a domain expert), CAPRI first performs a *lifting operation* over *D* to permit direct inference of complex selectivity relations over a joint representation, which involve input events as well as the hypotheses. Lifting operation evaluates each input CNF formula – for all input hypotheses in Φ – and outputs a lifted matrix *D*(Φ) to be processed further as in step 1. As an example, consider hypothesis *a b* ▹ *c* lifted input matrix *D* is:

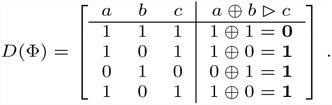

Note that the first row (profile {*a, b, c*}) contradicts the hypothesis, while all other rows support it.

#### Selectivity Topology (steps 3, 4, 5)

We exploit a compositional approach to test CNF hypotheses as follows: the disjunctive relations are grouped, and treated as if they were individual objects in *G*. For example, when a formula *φ* ▹ *d* where *φ* = (*a* ⋁*b*) ⋀*c* is considered, we assess *φ* ▹ *d* as whether (*a* ⋁*b*) ▹ *d* and *c* ▹ *d* hold – with the proviso that we treat (*a* ⋁*b*) as an individual event. Formally, with clauses (*φ*) we denote the disjunctive clauses in a CNF formula.

Nodes in the reconstruction are all input events together with all the disjunctive clauses of each input formula *φ*.

Edges in the reconstructed DAG are patterns that satisfy both conditions (1) and (2) of the selectivity relation ▹. Formally, CAPRI includes an edge between two nodes *φ* and *j* only if both Γ_*φ,j*_ = 𝒫 (*φ*) − 𝒫 (*j*) and 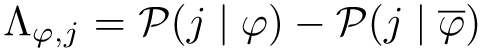 are strictly positive. Note that *φ* can be both a disjunctive clause as well as a singleton event. A function *π*(·) assigns a parent to each node that is not an input formula. Note that this approach works efficiently by nature of the lifted representation of *D*. The reconstructed DAG contains all the true positive patterns, with respect to ▹, plus spurious instances of ▹ which CAPRI subsequently removes in step 6 (cfr., the Supplementary Material for a proof of this statement).

Note that 𝒟 can be readily interpreted as a probabilistic graphical model, once it is augmented with a labeling function *α* : *N* → [0, 1], where *N* is the set of nodes – i.e., the genetic alterations – such that *α*(*i*) is the *independent probability* of observing mutation *i* in a sample, whenever *all of its parent* mutations (i.e., *π*(*i*)) are observed (if any). Thus 𝒟 induces a *distribution* of observing a subset of events in a set of samples (i.e., a probability of observing a certain *mutational profile* in a patient).

#### Maximum Likelihood Fit (step 6)

As the selectivity relation provides only a *necessary* condition, we must filter out all of its *spurious instances* that might have been included in 𝒟 (i.e., the possible *false positives*).

For any selectivity structure, spurious claims contribute to a reduction in the *likelihood-fit* relative to true patterns. Thus, a standard maximum-likelihood fit can be used to select and prune the selectivity DAG (including a *regularization term* to avoid over-fitting^10^). Here, we adopt the *Bayesian Information Criterion* (BIC), which implements *Occam’s razor* by combining log-likelihood fit with a *penalty criterion* proportional to the log of the DAG size via *Schwarz Information Criterion* (see [27]). The BIC score is defined as follows.

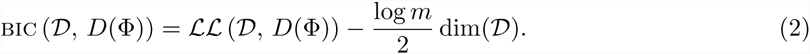

Here, *D*(Φ) is the lifted input matrix, *m* denotes the number of samples and dim(𝒟) is the number of parameters in the model 𝒟. Because, in general, dim(·) depends on the number of parents each node has, it is a good metric for model complexity. Moreover, since each edge added to 𝒟 increases model complexity, the regularization term based on dim(·) favors graphs with fewer edges and, more specifically, fewer parents for each node.

##### Algorithm 1 *CAncer PRogression Inference* (CAPRI)

1. **Input:** A set of events *G* = {*g*_1_*, …, g*_*n*_}, a matrix *D* ϵ {0, 1} ^*m*×*n*^ and *k* CNF causal claims Φ = {*φ*_1_ ▹ *e*_1_*, …, φ*_*k*_ ▹ *e*_*k*_ where, for any *i*, *ei φφsi* and *eiϵssG*;
2. [*Lifting*] Define the *lifting of D to D*(Φ) as the augmented matrix

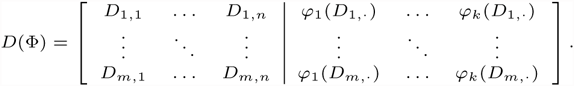

by adding a column for each *φi* ▹ *ci ∈* Φ, with *φi* evaluated row-by-row. Define then the coefficients Γ*i,j* = *P*(*i*) *- P*(*j*) and pairwise over *D*(Φ)
3. [*DAG nodes*] Define the set of nodes *N* = *G ∪*(∪_φ_clauses(φi)) which contains both input events and the disjunctive clauses in every input formula of Φ.
4. [*DAG edges*] Define a parent function *π* where *π*(*j* ∉ *G*) = ø – avoid edges incoming in a formula ^11^– and

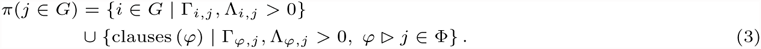

Set the DAG tos 𝒟(*N, π*).
5. [*DAG labeling*] Define the labeling *α* as follows

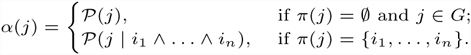
6. [*Likelihood fit*] Filter out all spurious causes from 𝒟by 𝒟 likelihood fit with the regularization BIC score and set *α*(*j*) = 0 for each removed edge.
7. 7:**Output:** the DAG 𝒟 and *α*;

At the end of this step,𝒟 and the labeling function are modified accordingly, based on the result of BIC regularization. By collecting all the incoming edges in a node it is possible to extract the patterns, which have been selected by CAPRI as the positive ones.

#### Inference Confidence: Bootstrap and Statistical Testing

To infer confidence intervals of the selectivity relations ▹, CAPRI employs *bootstrap with rejection resampling* as follows, by estimating a distribution of the marginal and joint probabilities. For each event, (*i*) CAPRI samples with repetitions rows from the input matrix *D* (bootstrapped dataset), (*ii*) CAPRI next estimates the distributions from the observed probabilities, and finally, (*iii*) CAPRI rejects values which do not satisfy 0 < 𝒫 (*i*) < 1 and 𝒫 (*i* | *j*) < 1 ∨ 𝒫 (*j i*) < 1, and iterates restarting from (*i*). We stop when we have, for each distribution, at least *K* values (in our case *K* = 100). Any inequality (i.e., checking temporal priority and probability raising) is estimated using the non-parametric Mann-Whitney U test^12^ with *p*-values set to 0.05. We compute confidence *p*-values for both temporal priority and probability raising using this test, which need not assume Gaussian distributions for the populations.

Once a DAG is inferred both *parametric and non-parametric bootstrapping methods* can be used to assign a confidence level to its respective pattern and to the overall model. Essentially, these tests consist of using the reconstructed model (in the parametric case), or the probabilities observed in the dataset (in the non-parametric case) to generate new synthetic datasets, which are then reused to reconstruct the progressions (see, e.g., [35] for an overview of these methods). The confidence is estimated by the number of times the DAG or any instance of ▹ is reconstructed from the generated data.

#### Complexity, Correctness and Expressivity

CAPRI has the following asymptotic complexity (Theorem 1, SI Section 2):

i. Without input hypotheses the execution is self-contained and polynomial in the size of *D*.
ii. In addition to the above cost, CAPRI tests input hypotheses of Φ at a polynomial cost in the size of ∣Φ∣. In this case, however, its complexity may range over many orders of magnitude depending on the structural complexity of the input set Φ consisting of hypotheses.

An empirical analysis of the execution time of CAPRI and the competing techniques on synthetic datasets is provided in the SI, Section 3.5.

CAPRI is a *sound and complete* algorithm, and its expressivity in terms of the inferred patterns is proportional to the hypothesis set Φ which, in turn, determines the complexity of the algorithm. With a proper set of input hypothesis, CAPRI can infer all (and only) the true patterns from the data, filtering out all the spurious ones (Theorem 2, SI Section 2).Without hypotheses, besides singleton and co-occurrence, no other patterns can be inferred (see Figure 2). Also, some of these claims might be spurious in general for more complex (and unverified) CNF formula (Theorem 3, SI Section 2).

## 4 Results and Discussion

To determine CAPRI’s relative accuracy (true-positives and false-negatives) and performance compared to the state-of-the-art techniques for *network inference*, we performed extensive *simulation experiments*. From a list of potential competitors of CAPRI, we selected: *Incremental Association Markov Blanket* (IAMB, [26]), the *PC algorithm* (see [25]), *Bayesian Information Criterion* (BIC, [27]), *Bayesian Dirichlet with likelihood equivalence* (BDE, [28]) *Conjunctive Bayesian Networks* (CBN, [8]) and *Cancer Progression Inference with Single Edges* (CAPRESE, [9]). These algorithms constitute a rich landscape of structural methods (IAMB and PC), likelihood scores (BIC and BDE) and hybrid approaches (CBN and CAPRESE).

Also, we applied CAPRI to the analysis of an atypical Chronic Myeloid Leukemia dataset of somatic mutations with data based on [16].

### 4.1 Synthetic data

We performed extensive tests on a large number of *synthetic datasets* generated by randomly parametrized progression models with distinct key features, such as the presence/absence of: (1) *branches*, (2) *confluences with patterns of co-occurrence*, (3) *independent progressions* (i.e., composed of disjoint sub-models involving distinct sets of events). Accordingly, we distinguish four classes of generative models with increasing complexity and the following features:

|  | *trees* | *forests* | *connected DAGs* | *disconnected DAGs* |
| --- | --- | --- | --- | --- |
| (1) | ✓ | ✓ | ✓ | ✓ |
| (2) | ✗ | ✗ | ✓ | ✓ |
| (3) | ✗ | ✓ | ✗ | ✓ |

The choice of these different type of topologies is not a mere technical exercise, but rather it is motivated, in our application of primary interest, by *heterogeneity of cancer cell types* and *possibility of multiple cells of origin*.

**Figure 4:**
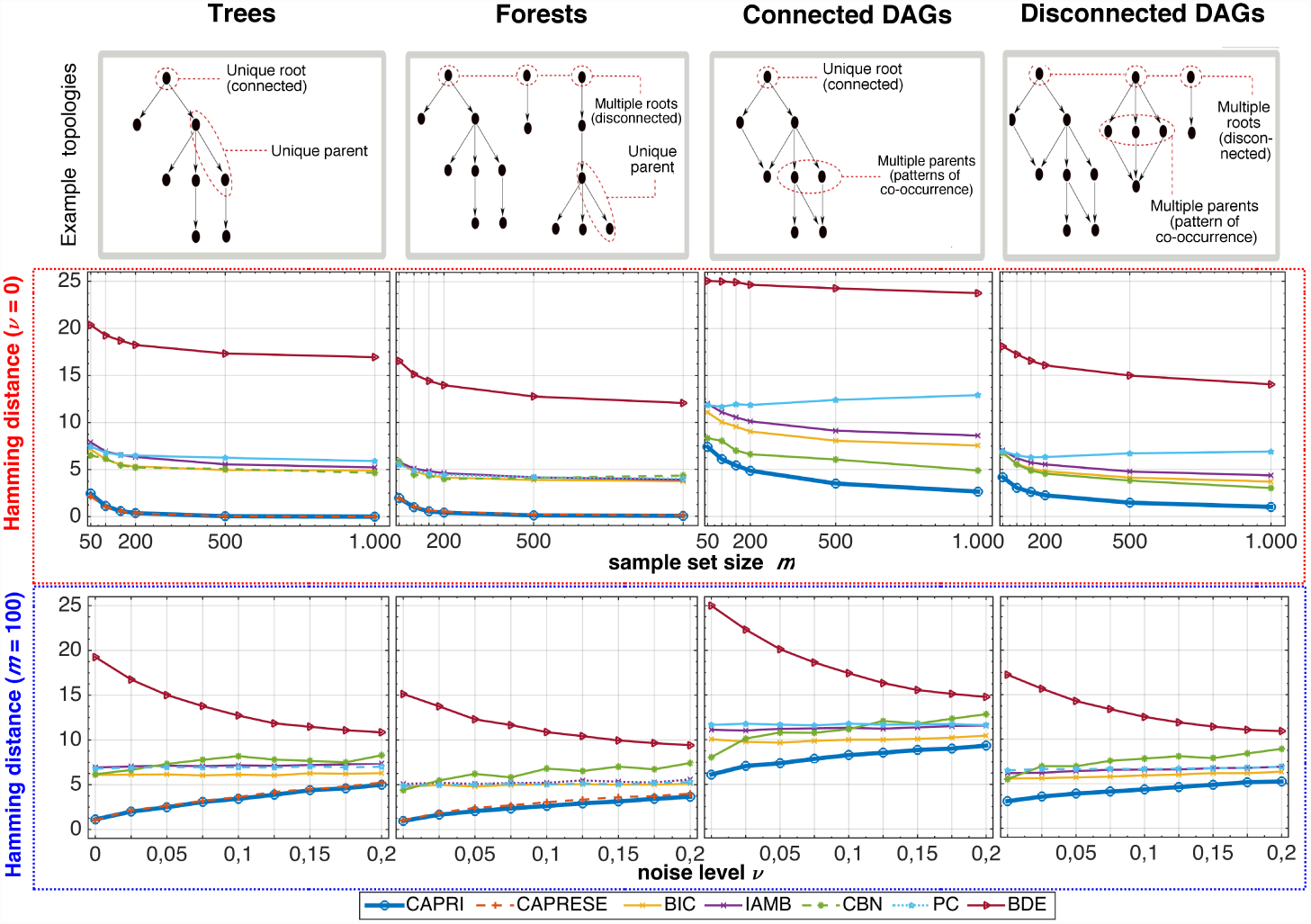
Comparative Study. Performance and accuracy of CAPRI (unsupervised execution) and other algorithms, IAMB, PC, BIC, BDE, CBN and CAPRESE, were compared using synthetic datasets sampled by a large number of randomly parametrized progression models–*trees*, *forests*, *connected* and *disconnected DAGs*, which capture different aspects of confluent, branched and het-erogenous cancer progressions. For each of those, 100 models with *n* = 10 events were created and 10 distinct datasets were sampled by each model. Datasets vary by number of samples (*m*) and level of noise in the data (*ν*) – see the Supplementary Information file for details. (*Red box*) Average *Hamming distance* (HD) – with 1000 runs – between the reconstructed and the generative model, as a function of dataset size (*mϵs* 50, 100, 150, 200, 500, 1000), when data contain no noise (*ν* = 0). The lower the HD, the smaller is the total rate of mis-inferred selectivity relations among events. (*Blue box*) The same is shown for a fixed sample set size *m* = 100 as a function of noise level in the data (*ν* ϵ{ 0, 0.025, 0.05, …, 0.2 }) so as to account for input *false positives* and *negatives*. See SI Section 3 for more extensive results on precision and recall scores and also including additional combinations of noise and samples as well as experimental settings.

To account for *biological noise* and *experimental errors* in the data we introduce a parameter *ν* ϵ (0, 1) which represents the probability of each entry to be random in *D*, thus representing a *false positive* (*ϵ*_+_) and a *false negative* rate (ϵ_−_): ϵ_+_ = ϵ_−_ = *ν/*2. The noise level complicates the inference problem, since samples generated from such topologies will likely contain sets of mutations that are correlated but causally irrelevant.

To have reliable statistics in all the tests, 100 distinct progression models per topology are generated and, for each model, for every chosen combination of sample set size *m* and noise rate *ν*, 10 different datasets are sampled (see SI Section 3 for our synthetic data generation methods).

Algorithmic performance was evaluated using the metrics *Hamming distance* (HD), *precision* and *recall*, as a function of dataset size, *ϵ*_+_ and *ϵ*_−_. HD measures the *structural similarity* among the reconstructed progression and the generative model in terms of the minimum-cost sequence of node edit operations (inclusion and exclusion) that transforms the reconstructed topology into the generative one^13^. Precision and recall are defined as follows: *precision* = TP/(TP + FP) and *recall* = TP/(TP + FN), where TP are the *true positives* (number of correctly inferred true patterns), FP are the *false positives* (number of spurious patterns inferred) and FN are the *false negatives* (number of true patterns that are *not* inferred). The closer both precision and recall are to 1, the better.

In Figure 4 we show the performance of CAPRI and of the competing techniques, in terms of Hamming distance, on datasets generated from models with 10 events and all the four different topologies. In particular, we show the performance: (*i*) in the case of noise-free datasets, i.e., *ν* = 0 and different values of the sample set size *m* and (*ii*) in the case of a fixed sample set size, *m* = 100 (size that is likely to be found in currently available cancer databases, such as TCGA (cfr., [4])) and different values of the noise rate *ν*. As is evident from Figure 4 CAPRI outperforms all the competing techniques with respect to all the topologies and all the possible combinations of noise rate and sample set size, in terms of average Hamming distance (with the only exception of CAPRESE in the case of tree and forests, which displays a behavior closer to CAPRI’s). The analyses on precision and recall display consistent results (SI Section 3). In other words, we demonstrate on the basis of extensive synthetic tests that CAPRI requires a much lower number of samples than the other techniques in order to converge to the real generative model and also that it is much more robust even in the presence of significant amount of noise in the data, irrespective of the underlying topology.

See SI Section 3 for a more complete description of the performance evaluation for all the analyzed combinations of parameters. There, we have shown that CAPRI is highly effective when the co-occurrence constraint on confluences is relaxed to *disjunctive* patterns, *even if no input hypotheses are provided*, i.e., Φ = ø. This result hints at CAPRI’s robustness to infer patterns with imperfect regularities. Finally, we also show that CAPRI is effective in inferring synthetic lethality relations in this case using the operator as introduced in Section 2, Approach; when a combination of mutations in two or more genes leads to cell death, while separately, the mutations are viable. In this case, candidate relations are directly input as Φ.

### 4.2 Atypical Chronic Myeloid Leukemia (aCML)

As a case study, we applied CAPRI to the mutational profiles of 64 aCML patients described in [16]. Through exome sequencing, the authors identify a recurring *missense point mutation* in the *SET-binding protein 1* (setbp1) gene as a novel aCML marker.

Among all the genes present in the dataset by Piazza *et al.*, we selected those either (*i*) mutated-considered any mutation type - in at least 5% of the input samples (3 patients), or (*ii*) hypothesised to be part of a functional aCML progression pattern in the literature ^14^. The input dataset with selected events is shown in Figure 5; notice that somatic mutations are categorised as *indel*, *missense point* and *nonsense point* as in [16]. In Figure 5 we show the model reconstructed by CAPRI (supervised mode, execution time 5 ≈ seconds) on this dataset, with confidence assessed via 1000 non-parametric bootstrap iterations. The model highlights several non trivial selectivity relations involving genomic events relevant to ACML development.

**Figure 5:**
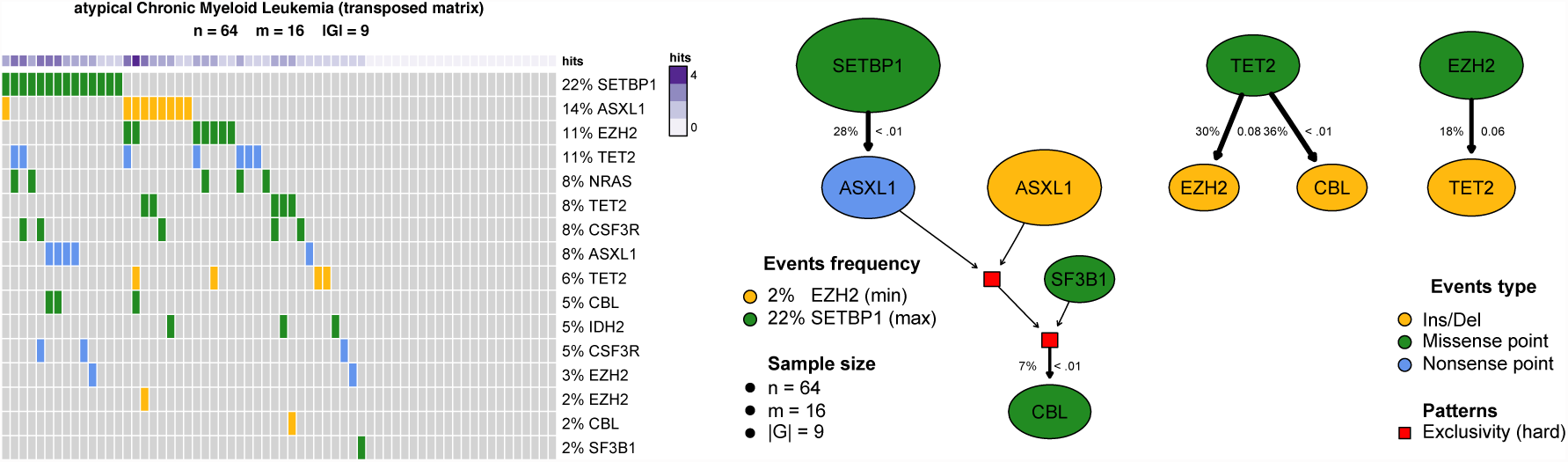
Atypical Chronic Myeloid Leukemia. *(left)* Mutational profiles of *n* = 64 aCML patients - exome sequencing in [16] - with alterations in *G* = 9 genes with either mutation frequency > 5% or belonging to an hypothesis inputed to CAPRI (SI Section 4). Mutation types are classified as *nonsense point*, *missense point* and *insertion/deletions*, yielding *m* = 16 input events. Purple annotations report the frequency of mutations per sample. *(right)* Progression model inferred by CAPRI in supervised mode. Node size is proportional to the marginal probability of each event, edge thickness to the confidence estimated with 1000 non-parametric bootstrap iterations (numbers shown leftmost of every edge). The p-value of the hypergeometric test is displayed too. Hard exclusivity patterns inputed to CAPRI are indicated as red squares. Events without inward/outward edges are not shown.

First, CAPRI predicts a progression involving mutations in setbp1, asxl1 and cbl, consistently with the recent study by [38], in which these genes were shown to be highly correlated and possibly functioning in a synergistic manner for aCML progression. Specifically, CAPRI predicts a selective advantage relation between missense point mutations in setbp1 and nonsense point mutations in asxl1. This is in line with recent evidence from [39] suggesting that setbp1 mutations are enriched among asxl1-mutated *myelodysplastic syndrome* (MDS) patients, and *in-vivo* experiments point to a driver role of setbp1 for that leukemic progression. Interestingly, our model seems also to suggest a different role of asxl1 *missense* and *nonsense* mutation types in the progression, yet more extensive studies (e.g., prospective or systems biology explanation) are needed to corroborate this hypothesis.

Among the hypotheses given as input to CAPRI, the algorithm seems to suggest that the exclusivity pattern among asxl1 and sf3b1 mutations selects for cbl missense point mutations. The role of the asxl1/sf3b1 exclusivity pattern is consistent with the study of [36] which shows that, on a cohort of 479 MDS patients, mutations in sf3b1are inversely related to asxl1 mutations.

Also, in [40] it was recently shown that asxl1 mutations, in patients with MDS, *myeloproliferative neoplasms* (MPN) and *acute myeloid leukemia*, most commonly occur as nonsense and insertion/deletion in a clustered region adjacent to the highly conserved *PHD* domain (see [41]) and that mutations of any type eventually result in a loss of asxl1 expression. This observation is consistent with the exclusivity pattern among asxl1 mutations in the reconstructed model, possibly suggesting alternative trajectories of somatic evolution for aCML (involving either asxl1 nonsense or indel mutations).

Finally, CAPRI predicts selective advantage relations among tet2 and ezh2 missense point and indel mutations. Even though the limited sample size does not allow to draw definitive conclusions on the ordering of such alterations, we can hypothesize that they may play a synergistic role in aCML progression. Indeed, [42] suggests that the concurrent loss of ezh2 and tet2 might cooperate in the pathogenesis of myelodysplastic disorders, by accelerating the overall tumor development, with respect to both MDSs and *overlap disorders* (MDS/MPN).

## 5 Conclusions

The *reconstruction of cancer progression models* is a pressing problem, as it promises to highlight important clues about the evolutionary dynamics of tumors and to help in better targeting therapy to the tumor (see e.g., [43]). In the absence of large longitudinal datasets, progression extraction algorithms rely primarily on *cross-sectional* input data, thus complicating the statistical inference problem.

In this paper we presented CAPRI, a new algorithm (and part of the TRONCO package) that attacks the progression model reconstruction problem by inferring *selectivity relationships* among “genetic events” and organizing them in a graphical model. The reconstruction algorithm draws its power from a combination of a scoring function (using Suppes’ conditions) and subsequent filtering and refining procedures, maximum-likelihood estimates and bootstrap iterations. We have shown that CAPRI outperforms a wide variety of state-of-the-art algorithms. We note that CAPRI performs especially well in the presence of noise in the data, and with limited sample size. Moreover we note that, unlike other approaches, CAPRI can reconstruct different types of confluent trajectories unaffected by the irregularities in the data – the only limitation being our ability to hypothesize these patterns in advance. We also note that CAPRI’s overall algorithmic complexity and convergence properties do offer several tradeoffs to the user.

Successful cancer progression extraction is complicated by tumor heterogeneity: many tumor types have molecular subtypes following different progression patterns. For this reason, it can be advantageous to cluster patient samples by their genetic subtype prior to applying CAPRI. Several tools have been developed that address this clustering problem (e.g., Network-based stratification [44] or COMET from [45]). A related problem is the classification of mutations into functional categories. In this paper, we have used genes with deleterious mutations as driving events. However, depending on other criteria, such as the level of homogeneity of the sample, the states of the progression can represent any set of discrete states at varying levels of abstraction. Examples include high-level hallmarks of cancer proposed by [46, 47], a set of affected pathways, a selection of driving genes, or a set of specific genomic aberrations such as genetic mutations at a more mechanistic level.

We are currently using CAPRI to conduct a number of studies on publicly available datasets (mostly from TCGA, [4]) in collaboration with colleagues from various institutions. In this work we have shown the results of the reconstruction on the aCML dataset published by [16], and in SI Section 4 we include a further example application on ovarian cancer ([48]), as well as a comparative study against the competing techniques. Furthermore, we are currently extending our pipeline in order to include pre-processing functionalities, such as patient clustering and categorization of mutations/genes into pathways (using databases such as the KEGG database (see [49]) and functionalities from tools like Network-based clustering, due to [44].

Encouraged by CAPRI’s ability to infer interesting relationships in a complex disease such as aCML, we expect that in the future CAPRI will help uncover relationships to aid our understanding sof cancer and eventually improve targeted therapy design.

## Acknowledgements

This research was funded by the NSF grants CCF-0836649 and CCF-0926166 and by Regione Lombardia (Italy) under the research projects RetroNet through the ASTIL Program [12-4-5148000-40]; U.A 053 and Network Enabled Drug Design project [ID14546A Rif SAL-7] Fondo Accordi Istituzionali 2009.

We also thank Francesca Ciccarelli, King’s College London, UK, and others for suggesting the “selectivity advantage” terminology. We would also like to thank all the participants of the Workshop and School on *Cancer, Systems and Complexity* held on Lake Como, Italy for many fruitful discussions there (csac.lakecomoschool.org). Finally, we are also indebted to Rocco Piazza, Universit`a degli Studi di Milano Bicocca, Italy, for all the data, insights and patience in explaining to us the biology of aCML.

## Footnotes

1 Unlike longitudinal studies, these cross-sectional data are derived from samples that are collected at unknown time points, and can be considered as “static”.

2 For instance, epigenetic alterations such as methylation and alterations in gene expression are not directly usable as input data for the algorithm. Notice that the selection of the relevant events is beyond the scope of this work and requires a further upstream pipeline, such as that provided, for instance, in [11, 3].

3 Suppes presents the relation in terms of causality; however, we avoid Suppes’ terminology as we build on just two of his many axioms, which only give rise to the notion of *prima-facie* causality.

4 Note that the conjunction ⋀ in our setting is interpreted differently from the classical notion (and the one adopted in e.g., [8]) since *a* ⋀ *b* ▹ *c* implies *a* ▹ *c* and *b* ▹ *c* in our framework. See also [17]. Moreover, note that the scope of this study is intentionally kept limited from further generalization of formulæ i.e., we will not consider statements of the form *φ*_*i*_ ▹ *φ*_*j*_, where the rightmost argument is a formula too.

5 A DAG is formed by a set of nodes and oriented edges connecting one node to another, such that there are no directed loops among them. See SI Section 1 for a technical definition.

6 This statement implies that there could be samples – i.e., patients – contradicting a pattern which still remains valid at a population level. For this reason a pattern *x* ⋀ *y* ▹ *z* is sometimes called a “noisy and”.

7 Theoretically, this assumption - common to other Bayesian learning problems - is necessary to prove CAPRI’s ability to extract the exact model in the optimal case of infinite samples. Practically, as all *relevant* events are hardly selectable a priori and sample size is finite, further statistics can be used to select the most relevant driver alterations – see also Section 4, Results and Discussion. Nonetheless, CAPRI can provide significant results even if this assumption is not or cannot be verified.

8 Formally, we require that *φ*_*i*_ ⋢ *e*_*i*_, where ⊑ represents the usual *syntactical* ordering relation among atomic events and formulas, and disallows for example *a* ⋁ *b* ▹ *a*.

9 In the current biomedical setting, the number of samples (*m*) is usually in the hundreds, while number of possible mutations (*n*) and hypotheses (*k*), absent any pre-processing, could be large, thus violating the assumption; in these cases, we rely on various commonly used pre-preprocessing filters to limit *n* to driver mutations, and *k* to simple hypotheses involving the driver mutations. However, in the future as the number of samples increases, we envision a more agnostic application.

10 In principle other regularisation strategies common to Bayesian learning could be used, e.g., Akaike information criterion (see [29] and references therein). In this paper, we prefer to work with BIC which, in general, trades model complexity to reduce false positives rate.

11 Although CAPRI is equipped with bootstrap testing it is still possible to encounter various degenerate situations. In particular, for some pair of events it could be that temporal priority cannot be satisfactorily resolved, i.e. there is no significant *p*-value for any edge orientation. Thus, loops might be present in the inferred prima facie topology. Nonetheless, some of these could be still disentangled by probability raising, while some might remain, albeit rarely. To remove such edges we suggest to proceed as follows: (*i*) sort these edges according to their *p*-value (considering both temporal priority and probability raising), (*ii*) scan the sorted list in decreasing order of confidence, (*iii*) remove an edge if it forms a loop.

12 The Mann-Whithney U test is a rank-based non-parametric statistical hypothesis test that can be used as an alternative to the Student’s t-test and is particularly useful if data are not normally distributed.

13 This measure corresponds to the sum of false positives and false negative and, for a set of *n* events, is bounded above by *n*(*n* − 1) when the reconstructed topology contains all the false negatives and positives.

14 Two *hard exclusivity* patterns - i.e., mutual exclusivity with “xor” - were tested, involving the mutations of: (*i*) genes asxl1 and sf3b1 (see [36]), which is present in the inferred progression model in Figure 5, and (*ii*) genes tet2 and idh2 (see [37]). The syntax in which the patterns are expressed is in the SI, Section 4.

